# Interaction network of SARS-CoV-2 with host receptome through spike protein

**DOI:** 10.1101/2020.09.09.287508

**Authors:** Yunqing Gu, Jun Cao, Xinyu Zhang, Hai Gao, Yuyan Wang, Jia Wang, Jinlan Zhang, Guanghui Shen, Xiaoyi Jiang, Jie Yang, Xichen Zheng, Jianqing Xu, Cheng Cheng Zhang, Fei Lan, Di Qu, Yun Zhao, Guoliang Xu, Youhua Xie, Min Luo, Zhigang Lu

**Affiliations:** The Fifth People’s Hospital of Shanghai, the Shanghai Key Laboratory of Medical Epigenetics, the International Co-laboratory of Medical Epigenetics and Metabolism, Ministry of Science and Technology, Institutes of Biomedical Sciences, Fudan University, Shanghai 200032, China; Institute of Pediatrics of Children’s Hospital of Fudan University, Shanghai 201102, China; Zhongshan-Xuhui Hospital, Fudan Univeristy, Shanghai 200020, China; Key Laboratory of Medical Molecular Virology (MOE/MOH), School of Basic Medical Sciences, Shanghai Medical College, Fudan University, Shanghai 200032, China; State Key Laboratory of Molecular Biology, CAS Center for Excellence in Molecular Cell Science, Institute of Biochemistry and Cell Biology, Shanghai Institutes for Biological Sciences, Chinese Academy of Sciences, Shanghai 200031, China; Shanghai Public Health Clinical Center, Fudan University, Shanghai 200433, China; Department of Physiology, University of Texas Southwestern Medical Center, Dallas, TX 75390, USA

**Author notes:** Corresponding author. (Z.L.); (M.L.); (Y.X.). These authors contributed equally to this work (Yunqing Gu, Jun Cao, Xinyu Zhang, Hai Gao, Yuyan Wang, Jia Wang).

## Abstract

Host cellular receptors are key determinants of virus tropism and pathogenesis. Virus utilizes multiple receptors for attachment, entry, or specific host responses. However, other than ACE2, little is known about SARS-CoV-2 receptors. Furthermore, ACE2 cannot easily interpret the multi-organ tropisms of SARS-CoV-2 nor the clinical differences between SARS-CoV-2 and SARS-CoV. To identify host cell receptors involved in SARS-CoV-2 interactions, we performed genomic receptor profiling to screen almost all human membrane proteins, with SARS-CoV-2 capsid spike (S) protein as the target. Twelve receptors were identified, including ACE2. Most receptors bind at least two domains on S protein, the receptor-binding-domain (RBD) and the N-terminal-domain (NTD), suggesting both are critical for virus-host interaction. Ectopic expression of ASGR1 or KREMEN1 is sufficient to enable entry of SARS-CoV-2, but not SARS-CoV and MERS-CoV. Analyzing single-cell transcriptome profiles from COVID-19 patients revealed that virus susceptibility in airway epithelial ciliated and secretory cells and immune macrophages highly correlates with expression of ACE2, KREMEN1 and ASGR1 respectively, and <u>A</u>CE2/A<u>S</u>GR1/K<u>R</u>EMEN1 (ASK) together displayed a much better correlation than any individual receptor. Based on modeling of systemic SARS-CoV-2 host interactions through S receptors, we revealed ASK correlation with SARS-CoV-2 multi-organ tropism and provided potential explanations for various COVID-19 symptoms. Our study identified a panel of SARS-CoV-2 receptors with diverse binding properties, biological functions, and clinical correlations or implications, including ASGR1 and KREMEN1 as the alternative entry receptors, providing insights into critical interactions of SARS-CoV-2 with host, as well as a useful resource and potential drug targets for COVID-19 investigation.

## MAIN TEXT

The global outbreak of COVID-19 caused by SARS-CoV-2 severely threatens human health ^1,2^. SARS-CoV-2 is a member of the beta-coronavirus genus, closely related to severe acute respiratory syndrome coronavirus (SARS-CoV), and both viruses use ACE2 as an entry receptor ^3-5^. SARS-CoV-2 is more than a respiratory virus, with multi-organ tropisms and causing complicated symptoms ^2,6-8^. Host cellular receptors play key roles in determining virus tropism and pathogenesis. Viruses bind to multiple host receptors for viral attachment, cell entry, and diverse specific host responses, including inducing cytokine secretion, stimulation of the immune response, or alteration of virus budding and release ^9-12^. However, apart from ACE2, little is known about SARS-CoV-2 receptors. Additionally, ACE2 cannot fully interpret SARS-CoV-2 tropism. The virus was detected in tissues with few ACE2 expression, such as liver, brain and blood, and even in lung, only a small subset of cells express ACE2 ^13,14^. The primary infection sites and clinical manifestations of SARS-CoV-2 and SARS-CoV also differ much, suggesting the involvement of other receptor(s) in SARS-CoV-2 host interaction ^1,2,15-18^. Therefore, a comprehensive understanding of SARS-CoV-2 cellular receptors is required.

Identification of receptors from virus-susceptible cells is limited to membrane proteins on specific cell types. We previously investigated ligand-receptor interactions using a cell based method, in which receptor-expressing cells were incubated with a tagged ligand and then an anti-tag for labelling and detection ^19,20^. It closely reassembles ligand-receptor interaction that occurring under physiological condition, and is usually used to confirm specific interactions. Based on this method, we developed a high-throughput receptor profiling system covering nearly all human membrane proteins, and used it to identify SARS-CoV-2 cellular receptors. Given SARS-CoV-2 S protein is the major receptor binding protein on the virus capsid, we performed profilling using S protein as the target. 5054 human membrane protein-encoding genes (91.6% of predicted human membrane proteins) were expressed individually on human 293e cells, and their binding with the extracellular domain of S protein (S-ECD) was measured (Fig. 1a). Twelve membrane proteins were identified that specifically interact with the S-ECD (Fig. 1b, c and Extended Data Fig.1), including the previously reported ACE2 ^4,5^ and SARS-CoV-specific CLEC4M (L-SIGN) ^21^.

**Figure 1:**
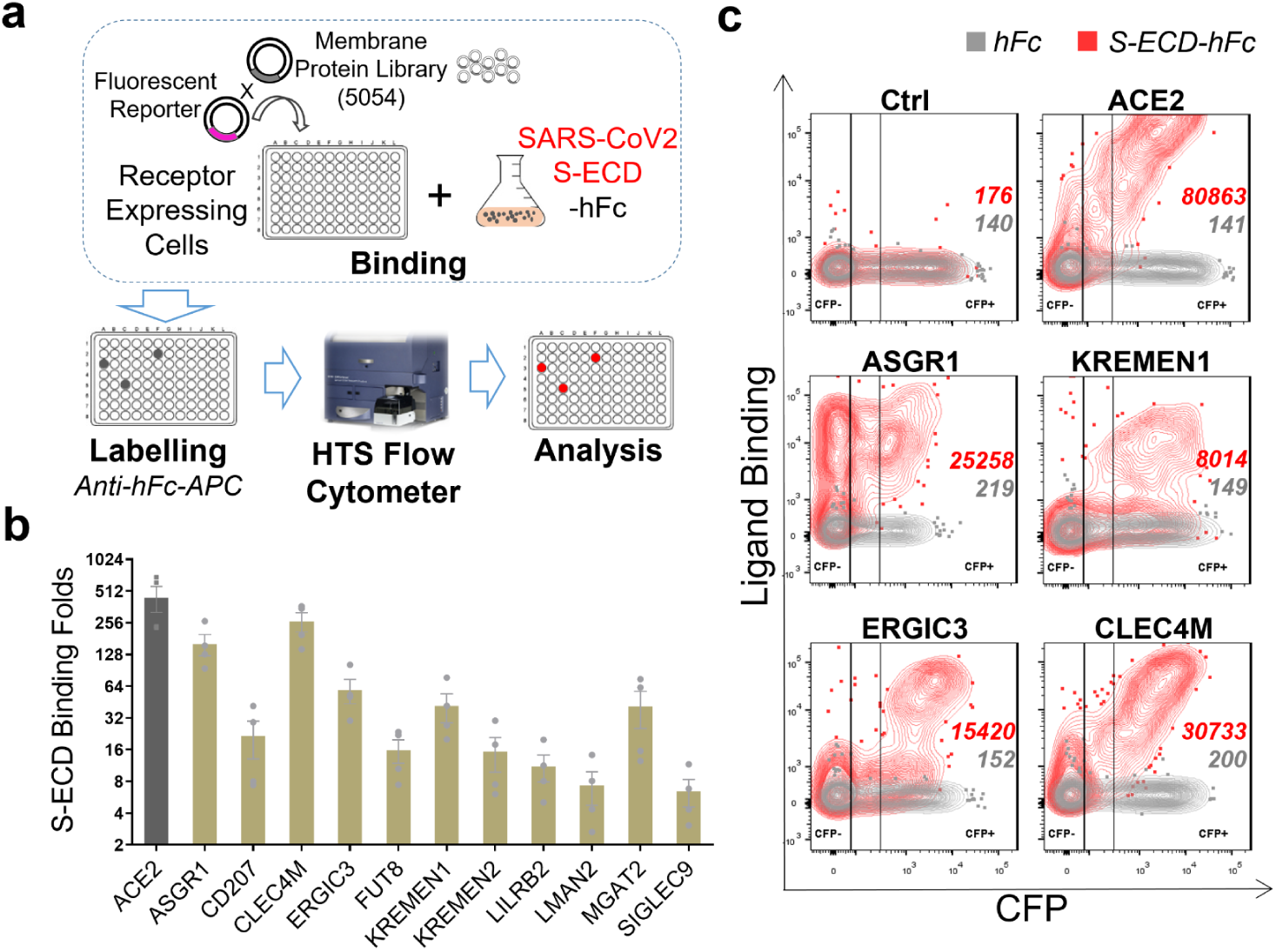
Genomic receptor profiling identifies twelve SARS-CoV-2 S binding receptors. **a**, Scheme of genomic receptor profiling. Plasmids encoding 5054 human membrane proteins were individually co-transfected with a CFP reporter into 293e cells. Cells were incubated with SARS-CoV-2 S-ECD-hFc protein, labelled using anti-hFc-APC antibody, binding was measured by flow cytometry. **b**, SIP identified S binding receptors. Relative binding of receptors with S-ECD-hFc compared to that with hFc control in CFP^+^ cells were shown. **c**, Representative flow dot plot showing the binding of S-ECD with top-ranking receptors.

The dissociation constants (Kd) of these interactions ranged from 12.4 - 525.4 nM (Table.1 and Extended Data Fig.2). ACE2 binds to S-ECD with a Kd of 12.4 nM, comparable to the previously reported Kd ^22^, and ACE2, CD207, CLEC4M, and KREMEN1 are all high-affinity receptors of the S protein, with comparable Kds. Binding domains on S protein were also examined, including the receptor binding domain (RBD), N-terminal domain (NTD) and S2 domain. The RBD and NTD are the major binding sites of S receptors. ACE2 only binds to the RBD, while CD207 and ERGIC3 bind exclusively with the NTD. The other receptors can bind to at least two domains, with CLEC4M, KREMEN1, and LILRB2 binding to all three domains, showing highest binding with NTD, RBD, and RBD, respectively (Table.1 and Extended Data Fig.3). Overall, these receptors showed diverse binding patterns, and they have a diverse range of biological functions and signaling properties (Extended Data Fig.4).

To determine whether these receptors can mediate virus entry independent of ACE-2, ACE2 was further knocked out in the HEK293T cell line (Extended Data Fig.5), which is low-sensitive to SARS-CoV-2 and SARS-CoV ^3,4,16^. We ectopically expressed the receptors in ACE2-KO 293T cells individually, and infected cells separately with pseudotyped SARS-CoV-2, SARS-CoV or MERS-CoV. KREMEN1-expressing cells showed clear evidence of SARS-CoV-2 infection, as did ASGR1, although to a lesser extent (Fig. 2a). Both receptors are specific to SARS-CoV-2, whereas ACE2 mediates the entry of both SARS-CoV-2 and SARS-CoV (Fig. 2a). ASGR1- and KREMEN1-dependent virus entry was confirmed with patient-derived SARS-CoV-2, with ASGR1 promoting higher levels of infection (Fig. 2b, c).

**Figure 2.**
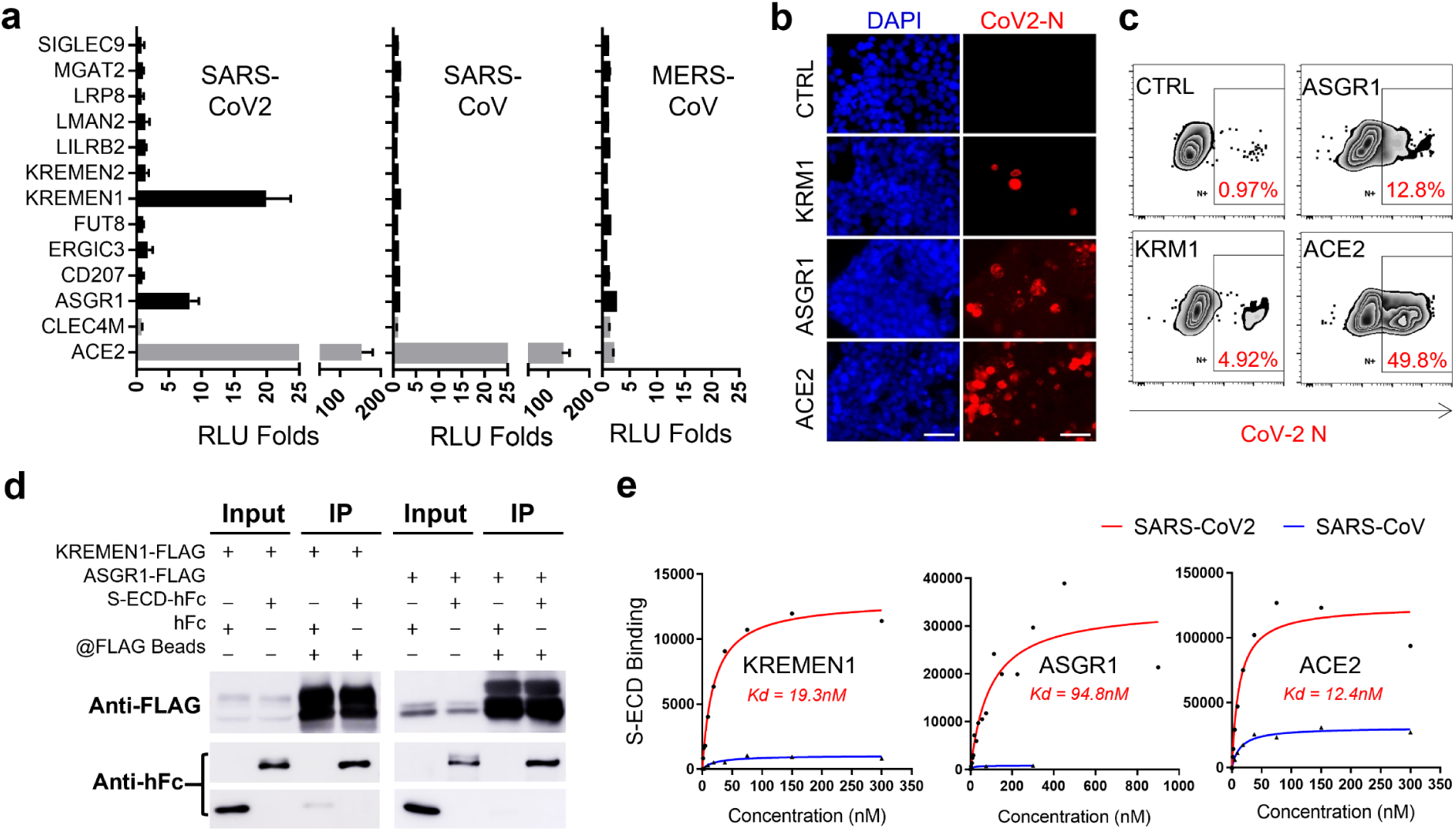
KREMEN1 and ASGR1 directly mediate SARS-CoV-2 entry. **a**, SIP-identified S-binding receptors were ectopically expressed in ACE2-KO 293T cells individually, followed by infection with pseudotyped SARS-CoV-2, SARS-CoV, and MERS-CoV separately. Luciferase activities relative to that of empty vector transfected cells were measured 60 hrs post infection. Data are presented as mean ± s.d (n=3). **b and c**, KREMEN1, ASGR1, or ACE2 transfected ACE2-KO 293T cells were infected with authentic SARS-CoV-2, and immune-fluorescence (B) or flow cytometry (C) were performed with antibody against the N protein of SARS-CoV-2 72hr post infection. Bar = 50μm. **d**, Co-immuno-precipitation was used to detect the interaction of S-ECD with full length KREMEN1 or ASGR1. **e**, KREMEN1, ASGR1 or ACE2 expressing 293e cells were incubated with different concentrations of S-ECD-hFc of SARS-CoV2 or SARS-CoV, separately, and S-ECD binding was monitored by flow cytometry to determine Kd.

Direct interaction of SARS-CoV-2 S protein with KREMEN1 and ASGR1 was confirmed by co-immuno-precipitation (Co-IP) (Fig. 2d). KREMEN1 and ASGR1 bind to S-ECD with Kds of 19.3 nM and 94.8 nM, respectively, the former Kd being comparable to that of ACE2 (12.4 nM). ACE2 has the highest maximum binding capacity for S-ECD, being ∼3- and ∼10-fold that of ASGR1 and KREMEN1, respectively (Fig. 2e), consistent with the SARS-CoV-2 sensitivities of cells expressing these receptors (Fig. 2a-c). Few binding of SARS-CoV S protein was observed with ASGR1 and KREMEN1 (Fig. 2e and Extended Data Fig.6), consistent with KREMEN1 and ASGR1 not mediating SARS-CoV infections (Fig. 2a). ACE2 binds exclusively to the RBD, KREMEN1 binds to all three domains of S-ECD, and with highest binding to the RBD, and ASGR1 binds to both the NTD and the RBD, the latter also with higher binding (Table. 1 and Extended Data Fig.3). Evidence indicates that the NTD is involved in entry coronavirus including SARS-CoV-2 ^23-26^, suggesting the potential importance of ASGR1 and KREMEN1 in SARS-CoV-2 infection. KREMEN1 is a high-affinity DKK1 receptor that antagonizes canonical WNT signaling ^27^, and is also the entry receptor for a major group of Enteroviruses ^28^. ASGR1 is an endocytic recycling receptor that plays a critical role in serum glycoprotein homeostasis ^29^ and has been reported to facilitate entry of Hepatitis C virus ^30^. Thus, ASGR1 and KREMEN1 directly mediate SARS-CoV-2 entry, together with ACE2, we refer to them as the ASK (<u>A</u>CE2/A<u>S</u>GR1/<u>K</u>REMEN1) entry receptors.

To investigate the clinical relevance of these entry receptors for SARS-CoV-2 susceptibility, we analyzed a recently published single cell sequencing (scRNA-seq) profile of the upper respiratory tract of 19 patients with COVID-19 ^31^. The dataset was derived from nasopharyngeal/pharyngeal swabs, and contains both the gene expression and virus infection status for individual cells, which are composed mainly of epithelial and immune populations. ACE2 is principally expressed in epithelial populations, as previously reported ^31^, whereas ASGR1 and KREMEN1 are enriched in both epithelial and immune populations (Fig. 3a). The majority of ASK^+^ cells express only one entry receptor (88.5%), and KREMEN1-expressing cells are the most abundant, being ∼5-fold more numerous than either ACE2- or ASGR1-expressing cells (Fig. 3b and Extended Data Fig.7a). SARS-CoV-2 is mainly observed in epithelial ciliated and secretory cells and immune non-resident macrophages (nrMa), which are also the major populations that express ASK receptors (Fig. 3b, c). Within SARS-CoV-2 positive cells (V^+^ cells), only 10.3% expressed ACE-2, suggesting other receptors will facilitate entry (Fig. 3c and Extended Data Fig.8).

**Figure 3.**
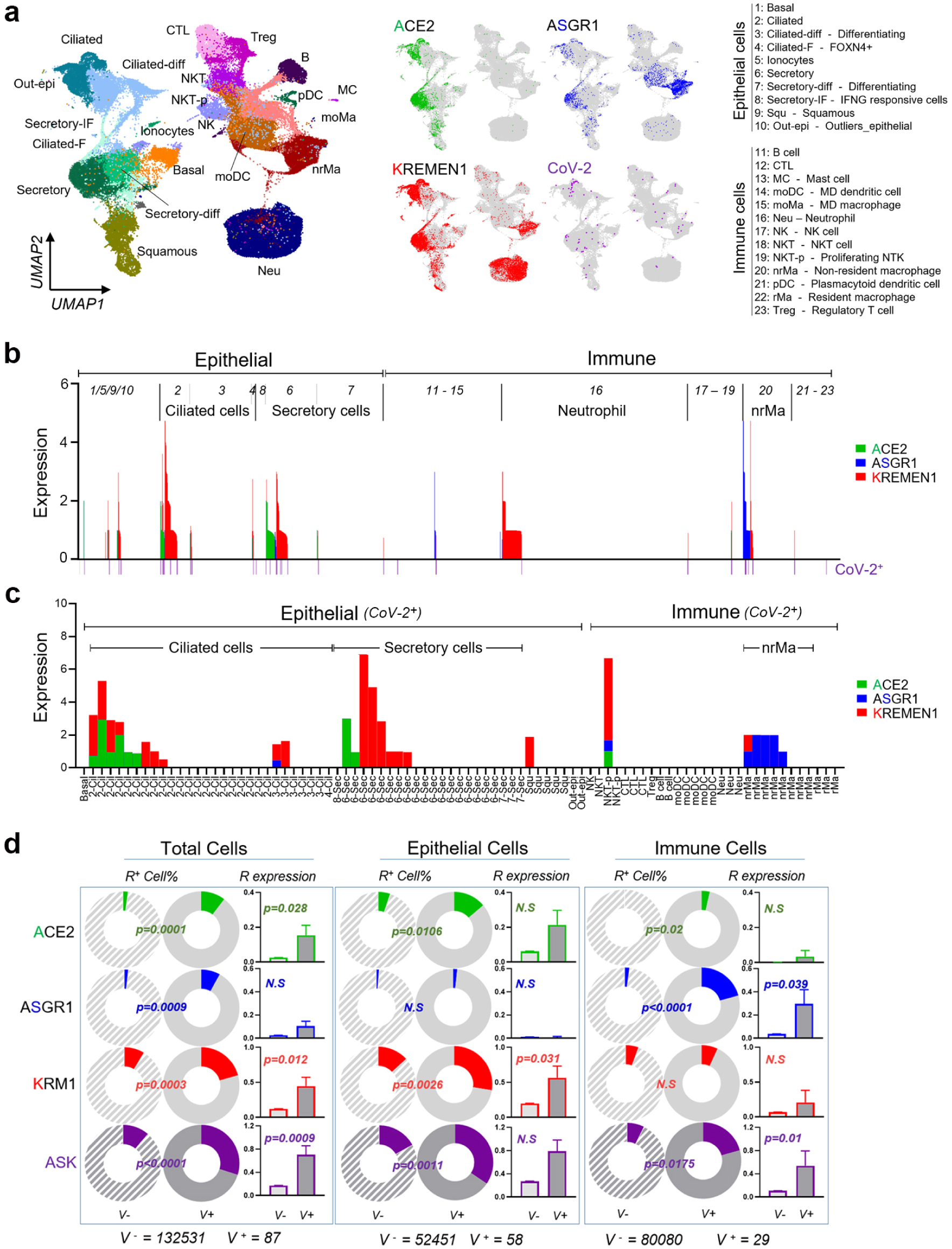
ASK entry receptors correlate significantly with SARS-CoV-2 susceptibility in the upper respiratory tract. **a**, Distribution of ACE2, ASGR1, KREMEN1 and SARS-CoV-2 in different cell populations of the upper airway. **b**, Overlapping map of ASK expression levels and virus infection pattern in different cell populations. **c**, ASK expression pattern in SARS-CoV-2 positive cells. **d**, Correlations of virus susceptibility with ASK receptors individually or in combination based on receptor positive cell percentage and receptor expression level.

We determined the correlation of the KREMEN1, ASGR1, and ACE2 entry receptors with SARS-CoV-2 susceptibility. In total cells, the receptor-positive cell percentage was significantly higher in V^+^ cells than in V^−^ cells for all three receptors (Fig. 3d). In epithelial populations, both ACE2 and KREMEN1 were substantially enriched in V^+^ cells, while in immune populations, only ASGR1 correlated with virus susceptibility, especially in macrophages (Fig. 3d and Extended Data Fig.7b). The epithelial ciliated and secretory cells are known target cells of SARS-CoV-2 ^14,31^. ACE2 displayed a more significant correlation with the virus susceptibility of ciliated cells when compared with KREMEN1, which was the only entry receptor that highly correlates virus susceptibility in secretory cells. Either in all cells or cell subpopulations, the ASK combination was usually more highly correlated with virus infection than individual receptors (Fig. 3d and Extended Data Fig.7b).

SARS-CoV-2 displays multi-organ tropism in COVID-19 patients ^6,12,13,17,32^. However, in virus-positive tissues, such as brain, liver, peripheral blood (PB) and even lung, ACE2 expression is few or only detected in a small subset of cells ^13,14,18^ (Extended Data Fig.9a, b), suggesting that ACE2 alone is difficult to explain the multi-organ tropisms of SARS-CoV-2. To determine whether ASK expression can predict tissue tropism better than ACE2 expression, we modeled a systemic host-SARS-CoV-2 interaction based on the expression of ASK entry receptors across human tissues (Fig. 4a). For a better comparison of different receptors from aspect of viral binding, the mRNA level was normalized with the S binding affinity of each receptor. ACE2 and ASGR1 are highly expressed in the gastrointestine and liver, respectively, while KREMEN1 is broadly expressed throughout the body. In virus-positive tissues, we found least one of the entry receptors is expressed (Extended Data Fig.9b and Fig. 4a). When ASK receptor expression levels were correlated with virus infection rates in different tissues reported in a recent biopsy study ^6^, the three receptors together correlated much better with virus susceptibility than any individual receptor (Fig. 4b). These results suggest that ASK expression underlies the multi-organ tropism of SARS-CoV-2, and is therefore can potentially predict viral tropism.

**Figure 4.**
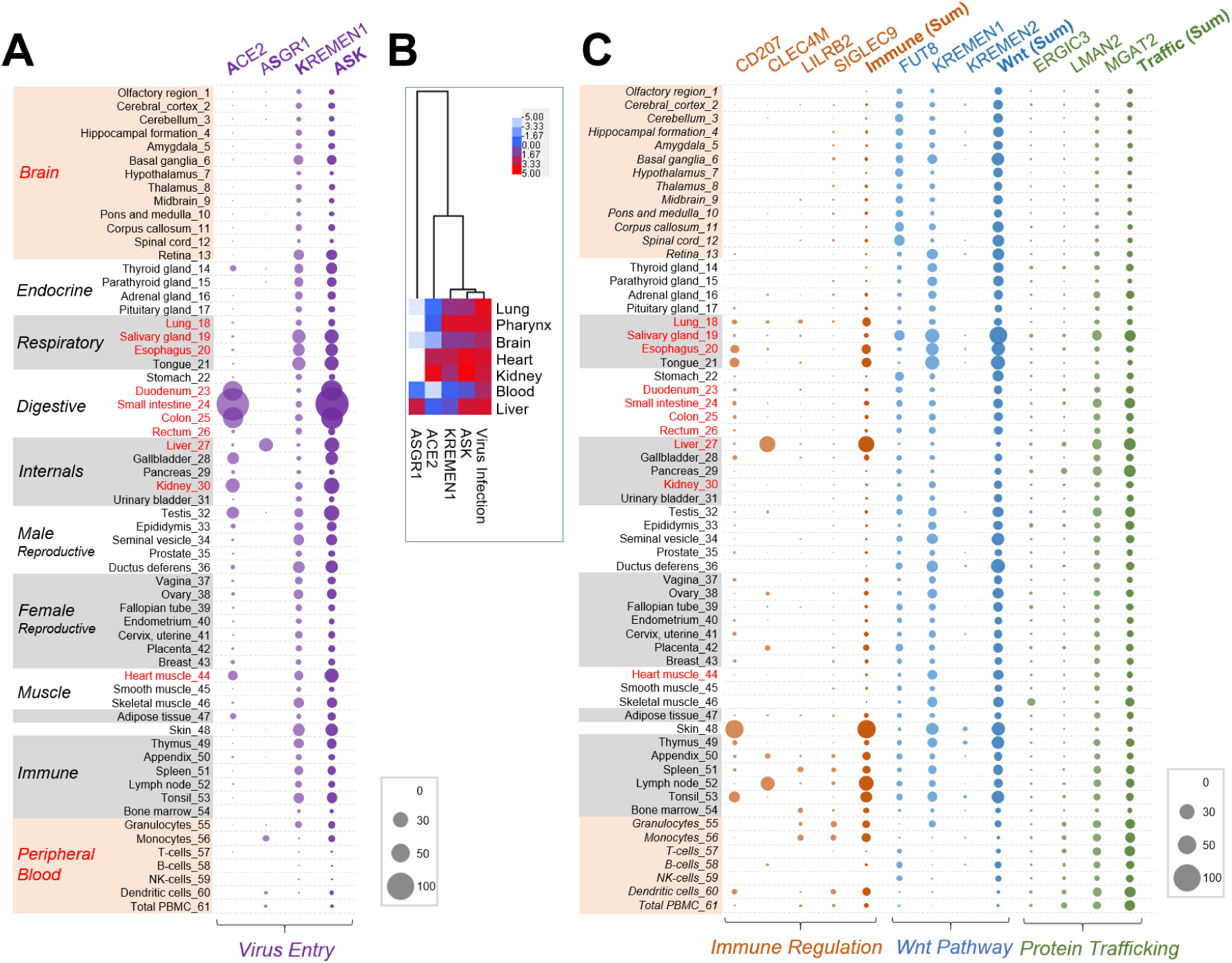
Systemic modeling of SARS-CoV-2 interactions within human tissues. The mRNA expression levels of each receptor in human tissues were derived from the human protein Atlas. Receptor-mediated SARS-CoV-2 binding potentials in each tissue were calculated by dividing receptor expression level with its affinity for S protein (Kd). **a**, Virus binding potentials of each tissue contributed by “entry group” ASK receptors. **b**, Virus infection rates (number of virus positive samples) in the indicated tissues were derived from a recent biopsy study ^6^, and these were clustered with ASK receptor-mediated virus binding potentials. **c**, Virus binding potentials of each tissue contributed by the S-binding receptors involved in immune regulation, the Wnt pathway and protein trafficking individually or in combination. In **a** and **c**, tissues or organs that were identified as positive to SARS-CoV-2 are labeled red.

**Table 1:** Characteristics of the interaction between receptors and SARS-CoV-2 S protein

| Gene Symbol | ACCN | MP Type | Kd Measurement |  | Relative Binding Folds |  |  |
| --- | --- | --- | --- | --- | --- | --- | --- |
|  |  |  | Kd (nM) | Std.Error | NTD | RBD | S2 |
| ACE2 | NM 021804 | I | 12.4 | 3.8 |  | **493** |  |
| ASGR1 | NM 001671 | II | 94.8 | 31.1 | 38.9 | **410** |  |
| CD207 | NM 015717 | II | 13.2 | 2.8 | **8.9** |  |  |
| CLEC4M | NM 014257 | II | 16.0 | 3.3 | **63.6** | 7.9 | 4.9 |
| ERGIC3 | NM 198398 | Multi | 525.4 | 40.5 | **16.4** |  |  |
| FUT8 | NM 004480 | II | 34.9 | 7.9 | **11.9** |  | 2.7 |
| KREMEN1 | NM 001039571 | I | 19.3 | 7.7 | 4.2 | **37.5** | 5.5 |
| KREMEN2 | NM 172229 | I | 60.0 | 14.2 |  | **14.0** |  |
| LILRB2 | NM 001080978 | I | 106.2 | 28.1 | 2.5 | **24.3** | 2.3 |
| LMAN2 | NM 006816 | I | 355.1 | 46.8 |  |  |  |
| MGAT2 | NM 002408 | II | 36.3 | 4.3 | **8.4** |  | 4.1 |
| SIGLEC9 | NM 014441 | I | 116.5 | 30.9 | 3.0 | **5.0** |  |

ACCN, Accession Number; MP Type, Membrane Protein Type.

Despite functioning in viral entry, virus-host receptor interactions could also induce cytokine secretion, apoptosis, and stimulation of the immune response, or alter virus budding and release ^9-12^. To gain insight into SARS-CoV-2 pathogenesis, we also modeled the host–SARS-CoV-2 interaction based on the tissue distribution of all the other receptors identified, which were classified according to their functions in immune regulation, the Wnt pathway, and protein trafficking. The interaction map revealed that expression of immune receptors is prominent in immune organs, as well as respiratory organs, and the liver (Extended Data Fig.9 and Fig. 4a), consistent with the respiratory manifestation and frequent liver injury in COVID-19 patients ^1,2,32,33^. Given that CD207, CLEC4M, LILRB2 and SIGLEC9 all are mainly expressed in myeloid cells (Extended Data Fig.10) and that COVID-19 is associated with hyperactivation of myeloid populations ^12,34,35^, it is possible that these receptors may drive monocyte and macrophage activation in COVID-19 and contribute to disease pathophysiology.

ERGIC3, LMAN2, and MGAT2, which are involved in protein trafficking, display an approximately similar expression levels across most human tissues (Extended Data Fig.9 and Fig. 4c). ERGIC3 and LMAN2 are the components of the endoplasmic reticulum-Golgi intermediate compartment (ERGIC), which is essential for coronavirus assembly and budding ^36 37^, while LMAN2 and ERGIC1 were recently found to interact specifically with nonstructural protein Nsp7 and Nsp10 of SARS-CoV-2 respectively ^38^. Whether and how they cooperate during virus life cycle are worth further investigation. Expression of receptors of the Wnt pathway group is prominent in salivary gland, tongue, esophagus, and brain (Extended Data Fig.9 and Fig. 4c). Wnt/β-catenin signaling is critical in taste bud cell renewal and behavioral taste perception ^39,40^ and KREMEN1/2 plus FUT8 are all negative regulators of this pathway ^27,41^. Loss of smell and taste has frequently been observed in COVID-19 patients ^42,43^, suggesting SARS-CoV-2 may act through these receptors to affect Wnt/β-catenin signaling and therefore taste loss.

The affinity-based interactions between SARS-CoV-2 and cellular receptors are key determinants of virus tropism and pathogenesis. Determining cells or tissues that express receptors should allow better characterization of the pathway for virus infection and help understand COVID-19 disease progression. Our genomic receptor profiling of most human membrane proteins has identified two additional virus entry receptors, ASGR1 and KREMEN1, independent of known ACE2. The combined ASK expression pattern predicts viral tropism much more closely than any individual entry receptor from cell to tissue levels. Our results also suggested that SARS-CoV-2 entry into different type of cells rely on different receptors, and ASK receptors underlie the tropism of SARS-CoV-2. Notably, ASGR1 and KREMEN1 do not mediate the entry of SARS-CoV, plausibly explaining the difference of these two viruses in primary infection sites and clinical manifestations. Unlike ACE2, which only binds to the RBD, ASGR1 and KREMEN1 bind to both the RBD and NTD. NTD is implicated in coronavirus entry ^25,26^, and several neutralizing antibodies from convalescent COVID-19 patients recognizes NTD ^23,24^, suggesting that the domain plays a role during SARS-CoV-2 infection, and that antibodies against the NTD may act through ASGR1 or KREMEN1.

The twelve SARS-CoV-2 receptors that bind S protein have diverse binding properties, functions, and tissue distributions. Integrating this panel of receptors with virological and clinical data should lead to the identification of infection and pathological mechanisms and targets. It is plausible that alternative binding receptors exert context-dependent regulatory effects, leading to differential signaling outcomes, ultimately influencing infection patterns, immune responses and clinical progression. Our study provides insight into critical virus-host interactions, tropisms, and pathogenesis of SARS-CoV-2, as well as potential targets for drug development against COVID-19.

## METHODS

### Ethics statement

All procedures in this study regarding authentic SARS-CoV-2 virus were performed in biosafety level 3 (BSL-3) facility, Medical School of Fudan University.

### Cell culture and transfection

Vero E6 cells and HEK293T cells were cultured in DMEM supplemented with 10% FBS at 37°C in 5% CO_2_ and the normal level of O_2_. HEK293e cells were cultured in serum-free FreeStyle 293 Medium (Invitrogen) with 120 rpm rotation at 37°C in 5% CO_2_ and the normal level of O_2_. For transient overexpression in 293T and 293e, plasmids were transfected using Lipofectamine 2000 (Invitrogen) according to the manufacture provided protocol.

### Genomic receptor profiling

To prepare SARS-CoV-2 S-ECD-hFc or control hFc containing condition medium, pCMV-S-ECD-hFc or pCMV-secreted-hFc plasmid was transfected into 293e cells, and condition medium was collected 4 days post transfection and filtered with 0.45um filter for screening. To prepare receptor expressing cells, plasmids encoding 5054 human membrane protein were individually co-transfected with CFP reporter vector (5:1) into 293e cells in 96 deep-well-plate. 2-5×10^4^ membrane protein expressing cells per well were collected 48hrs after transfection, washed once with PBS/2%FBS and incubated with 1ml SARS-CoV-2 S-ECD or hFc control condition medium for 1hr on ice. Supernatant was discarded after centrifugation and washed once with PBS/2%FBS, the cells were then labelled with Anti-hFc-APC (Jackson Lab) antibody for 20min, and washed once with PBS/2%FBS. The binding of S-ECD to the cells were measured by HTS flow cytometry (BD CantoII). The flow data were analyzed with FlowJo software. Relative binding of receptor (CFP^+^ cells) to S-ECD-hFc compared with that to hFc control were measured.

### Co-IP and Kd measurement

Receptor expressing cells were lysed with RIPA buffer (Sigma) and centrifuged for 15min at 15000rpm at 4°C, the cell lysate were collected. Purified hFc-tagged S-ECD proteins (final concentration of 10ug/ml) were added into cell lysate together with anti-FLAG beads, and incubated at 4°C for overnight. Beads were washed three times with the RIPA buffer, and the samples were prepared for western blot with anti-hFc or anti-FLAG antibodies. For Kd measurement, receptor coding plasmid was co-transfected with CFP reporter vector (5:1) into 293e cells. Cells were collected 48hr after transfection. ∼10^4^ cells per well were used for binding with series diluted purified S-ECD-hFc proteins as described in the experiment of Secretome Interaction Profiling. The flow data were analyzed with FlowJo software. Ligand binding value at each ligand concentration was normalized by equation of [(APC-MFI of CFP^+^)-(APC-MFI of CFP^−^)] - [(APC-MFI of CFP^+^)-(APC-MFI of CFP^−^)] _zero ligand concentration_. Kd and Bmax (maximum binding value) were calculated with Prism8 software.

### Protein purification and western blot

For purification of SARS-CoV-2 S-ECD-hFc, RBD-hFc, NTD-hFc, S2-hFc and SARS-CoV S-ECD, the plasmids were transfected into 293e cells, and condition medium was collected 4 days post transfection and filtered with 0.45um filter. hFc tagged proteins were purified using Protein A affinity column and then desalted to PBS solution with AKTA purifier system. Proteins were concentrated by 10KDa cutoff spin column (Amicon). For western blot, samples was separated by SDS-PAGE gel and transferred to nitrocellulose membrane. The membrane was labeled with the primary antibody and then HRP-conjugated secondary antibody at suggested concentration, and detected by ECL kit (Beyotime).

### ACE2 knockout 293T stable cell line

ACE2 small guide RNA was constructed into pSLQ1651 (Addgene #51024) *(44)* with a targeting sequence of CTTGGCCTGTTCCTCAATGGTGG. ACE2 sgRNA plasmid or Cas9Bsd plasmid (Addgene #68343) *(45)* were co-transfected with psPAX2 and pMD2G plasmids into 293T cells by Lipofectamine 2000 (Invitrogen) according to the manufacture provided protocol. Lentivirus were collected 72hr post transfection to infect 293T cells. ACE2 KO 293T stable cell line were obtained by single cell dilution.

### Pseudotyped coronavirus packaging and infection

For pesudotyped SARS-CoV-2, SARS-CoV and MERS-CoV, S protein encoding pCDNA3.1 plasmids were mixed with pNL4-3.Luc.R vector separately with a ratio of 1:1, and transfected into 293T cells using Lipofectamine 2000. Virus-containing supernatant was collected 48-72 hours post-transfection and filtered through 0.45um PES membrane filter (Millipore). For infection, cells were seeded into 96 well plate with ∼2×104 cells per well, 50ul virus-containing supernatant per well was added. Luciferase activities were measured 48hr post infection with Bright-Lumi™ Firefly Luciferase Reporter Gene Assay Kit (Beyotime, RG051M) and multifunctional microplate reader (TECAN 200pro).

### Authentic SARS-CoV-2 generation and infection

SARS-CoV-2/MT020880.1 were expanded in Vero E6 cells. Cells were collected 50hr post-infection and lysed by freeze-thaw method. Virus containing supernatants were collected by centrifugation at ∼2500xg for 10 minutes, and aliquot and stored at −80°C. For infection, targeted cells were incubated with fresh medium diluted virus supernatant at MOI of 0.1 for 48hrs. SARS-CoV-2 replication was examined by immuno-fluorescence and flow cytometry with anti SARS-CoV-2 N protein antibody.

### Data analysis and statistics

Gene Ontology Enrichment Analysis was performed by R bioconductor. For host-virus interaction map, receptor expression in each tissues were obtained from human Protein Atlas (https://www.proteinatlas.org/). mRNA expression level was normalized by dividing the expression level with the Kd of each receptor. Virus infection rates of tissues were obtained from the study published by Puelles et al. Cluster was performed with R package. For single cell sequencing (scRNA-seq) profile of the upper airway tract with COVID-19, the count, viral load and metadata are obtained from Magellan COVID-19 data explorer at https://digital.bihealth.org. Chi-square test and student’s t-test were performed to compare receptor percentage and receptor expression value in different cell populations respectively. All tests were two sided. P value <0.05 was designated significance.

## Reporting Summary

Further information on research design is available in the Nature Research Reporting Summary linked to this paper.

## Data availability

Receptor expression levels in each tissues were obtained from human Protein Atlas (https://www.proteinatlas.org/). Single cell sequencing (scRNA-seq) profile of the upper airway tract with COVID-19 and the metadata were obtained from Magellan COVID-19 data explorer at https://digital.bihealth.org. All data supporting the findings of this study are available within the paper or in the extended data.

## ACKNOWLEDGEMENTS

We thank Guy Riddihough of Life Science Editors for manuscript discussion and revision, and the staff in the Biosafety Level 3 Laboratory of Fudan University for experiments helping. This study was funded by the National Natural Science Foundation of China (projects 81873438 to Z.L., 81873922 to M.L.).

## AUTHOR CONTRIBUTIONS

Z.L., M.L., Y.X., and G.X. conceived the project. Y.G., X.Z. and J.C with the help from J.Z., X.J., J.W., J.Y., X.Z., W.Y., Y.Z., performed receptor profiling, characterizing receptor-ligand interaction. M.L. H.G. and Y.W. performed virus related experiments with the help from G.S., X.J., F.L.. Z.L., M.L., J.W. Y.G., J.C. H.G. and Y.W. performed bioinformatics analysis and analyzed the data. Z.L., M.L., Y.X., G.X. and Y.Z. wrote the manuscript.

## EXTENDED DATA FIGURES

**Extended Data Fig.1.**
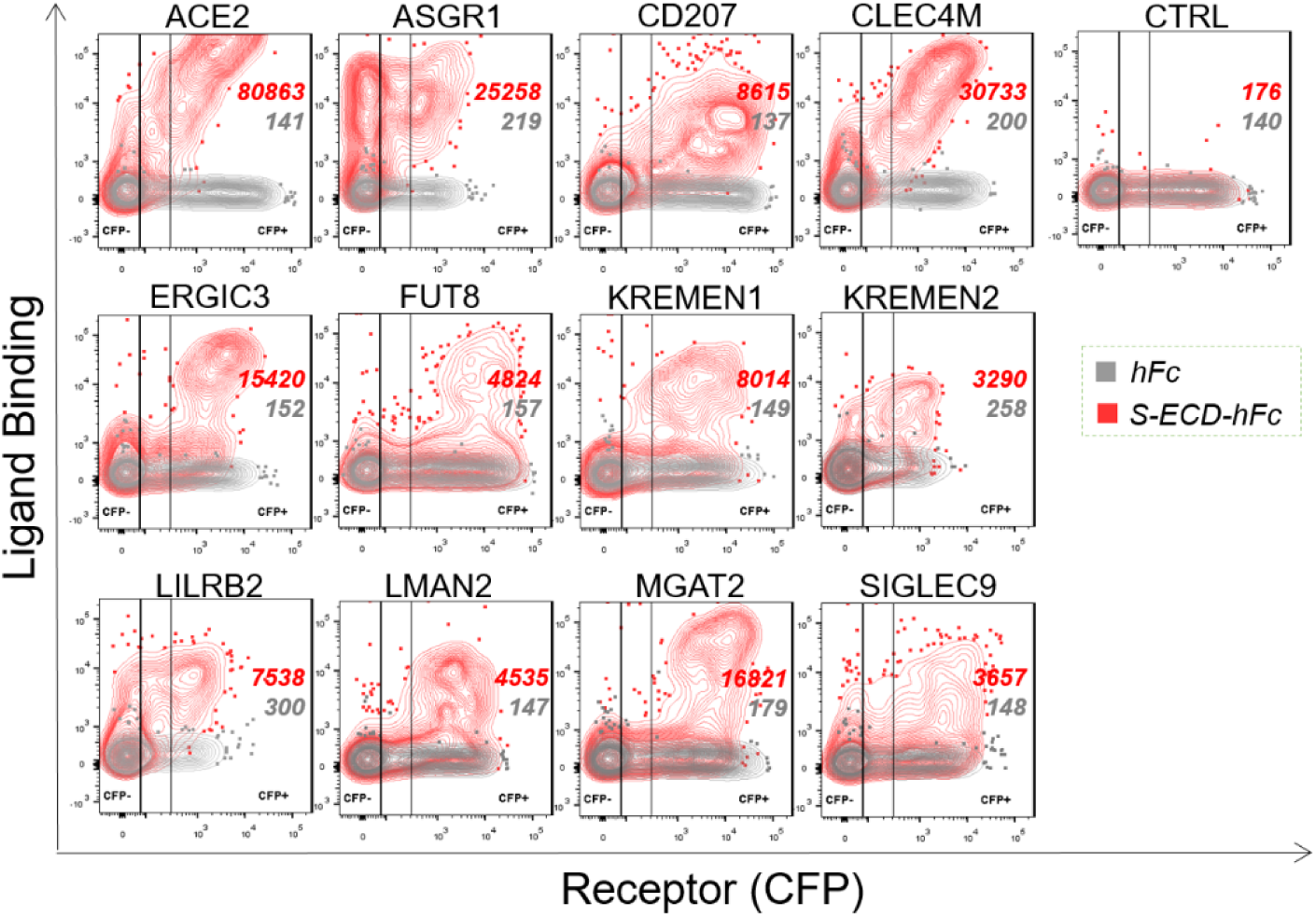
Binding of S-ECD with the profilling identified receptors. Plasmids encoding the indicated receptors were individually co-transfected with CFP reporter into 293e cells. The cells were incubated with SARS-CoV-2 S-ECD-hFc protein or hFc control protein, and then labelled by Anti-hFc-APC antibody, binding were measured by flow cytometry. Binding of S-ECD or hFc control to receptor were shown (Mean Fluorescent Intensity (MFI) of APC fluorescence).

**Extended Data Fig.2.**
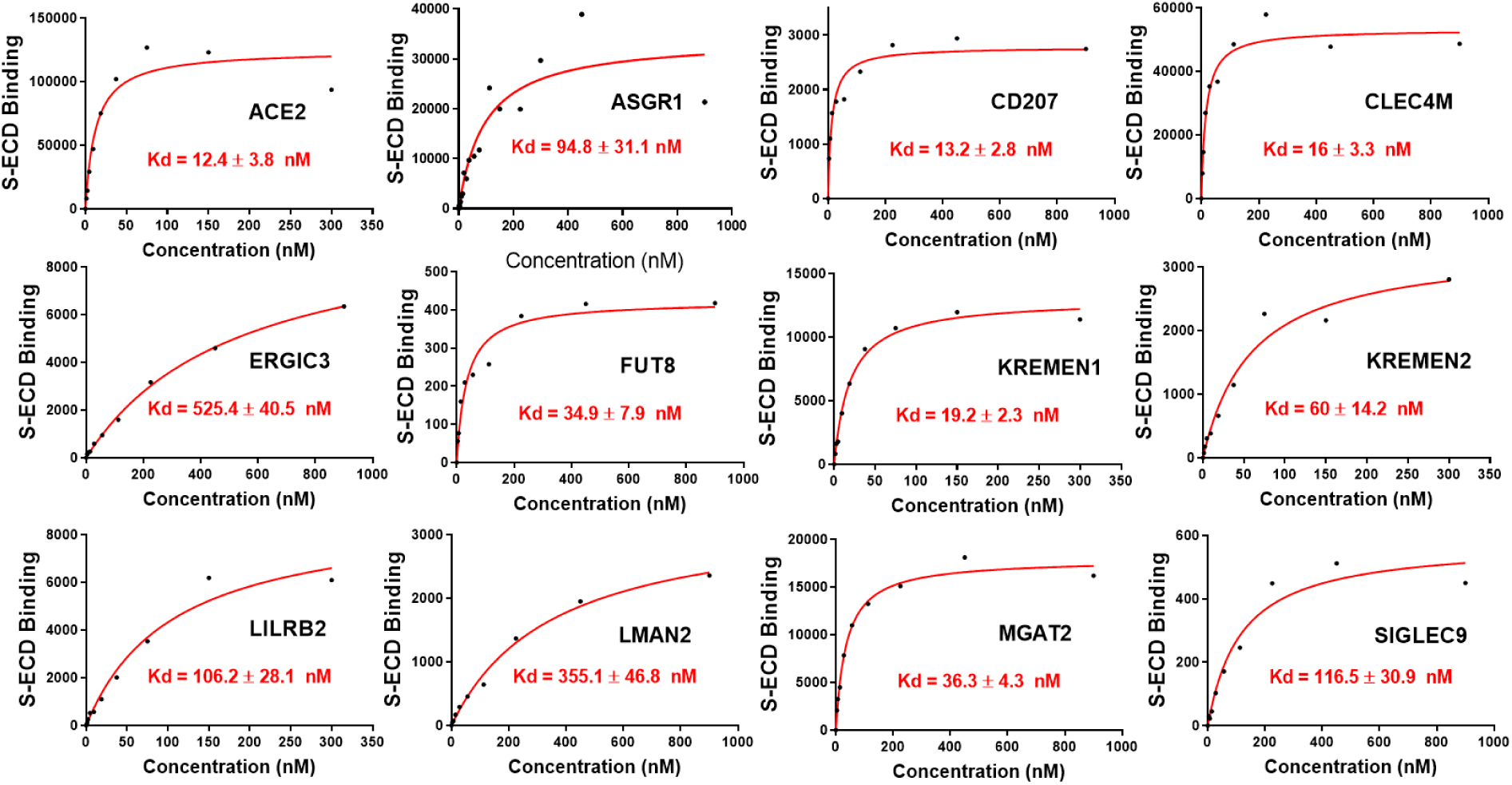
Kd measurement of the interaction of SARS-CoV-2 S-ECD with its receptors. 293e cells expressing the indicated receptors were incubated with serially diluted concentrations of SARS-CoV2 S-ECD-hFc, S-ECD binding were determined by flow cytometry for Kd measurement.

**Extended Data Fig.3.**
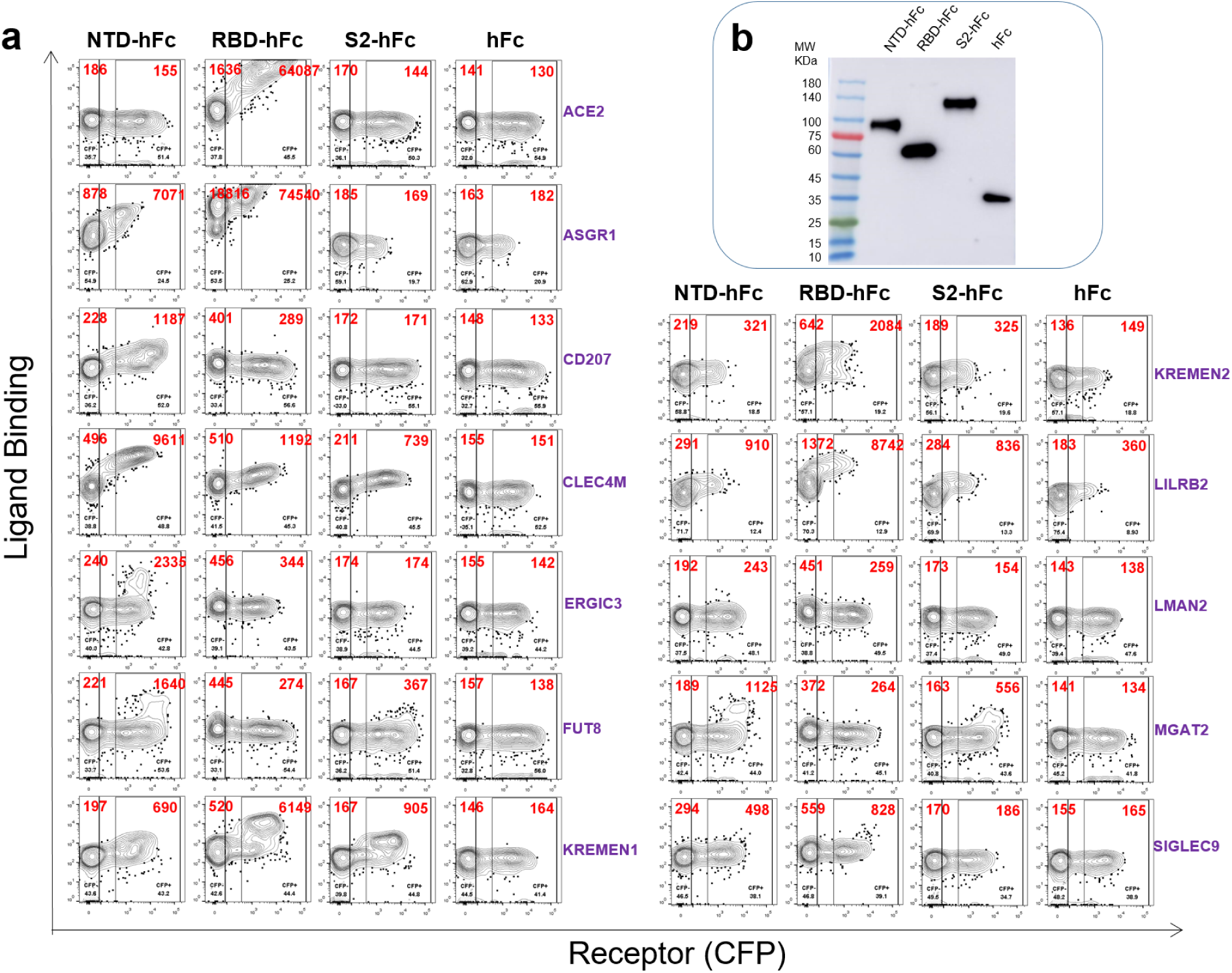
Analysis of binding domain on SARS-CoV-2 S protein. **a**, 293e cells expressing the indicated receptors were incubated with NTD-hFc, RBD-hFc, S2-hFc or hFc control separately. Binding were measured by flow cytometry. **b**, Western blot with anti-hFc antibody showing the ligand proteins used in this assay.

**Extended Data Fig.4.**
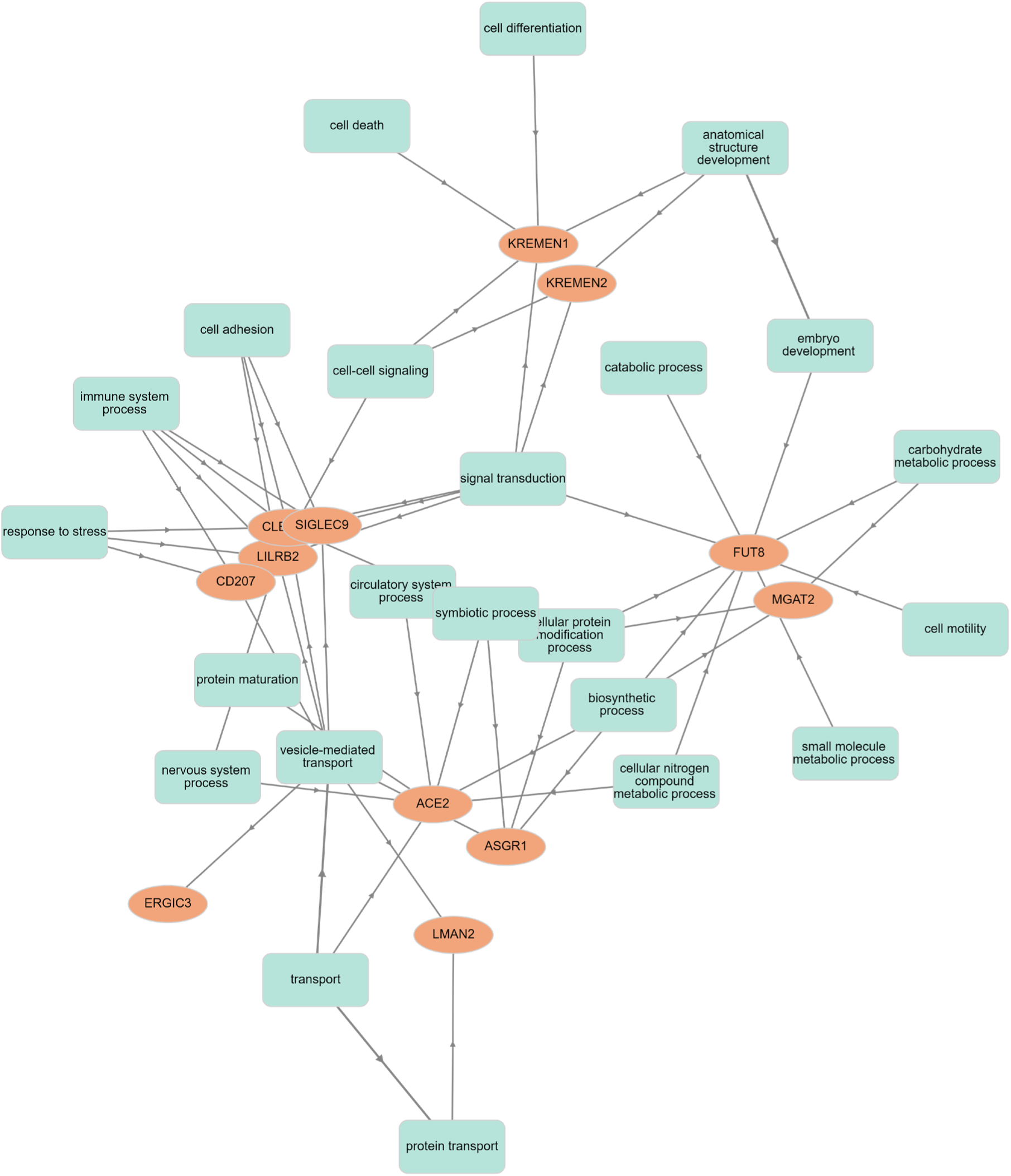
Biological function network of twelve SARS-CoV-2 S receptors.

**Extended Data Fig.5.**
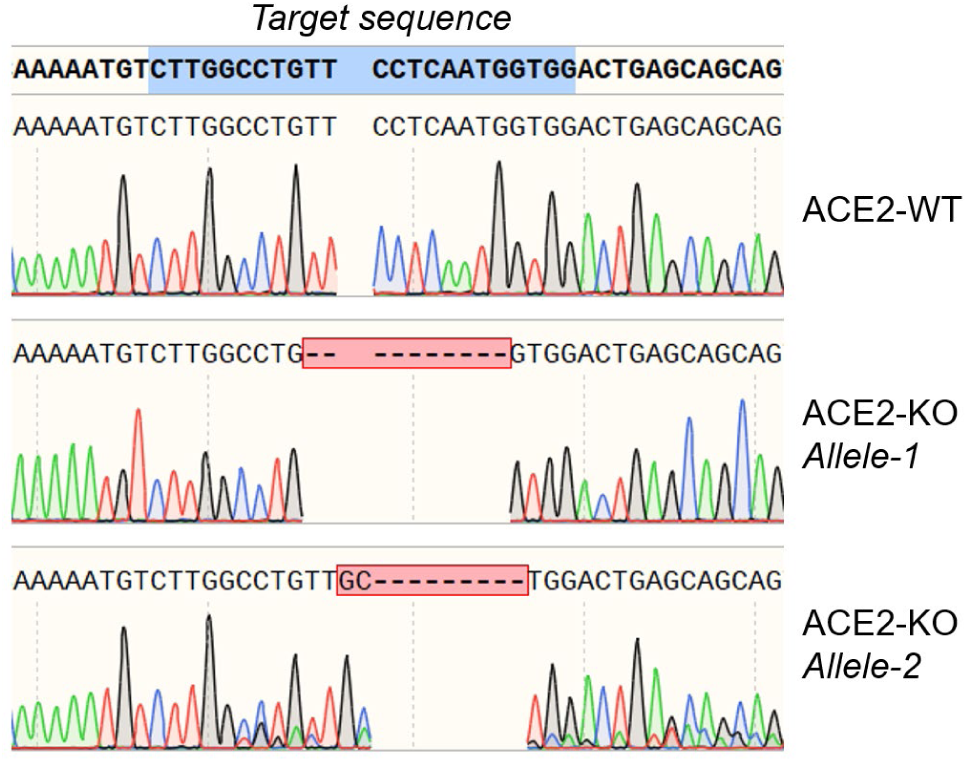
Genotyping of ACE2 KO 293T cell line. ACE2 exon1 was PCR amplified from ACE2-WT/KO 293T cells for sequencing. Gene editing at ACE2 locus on both alleles were shown. Both editing result in frame-shift of ACE2.

**Extended Data Fig.6.**
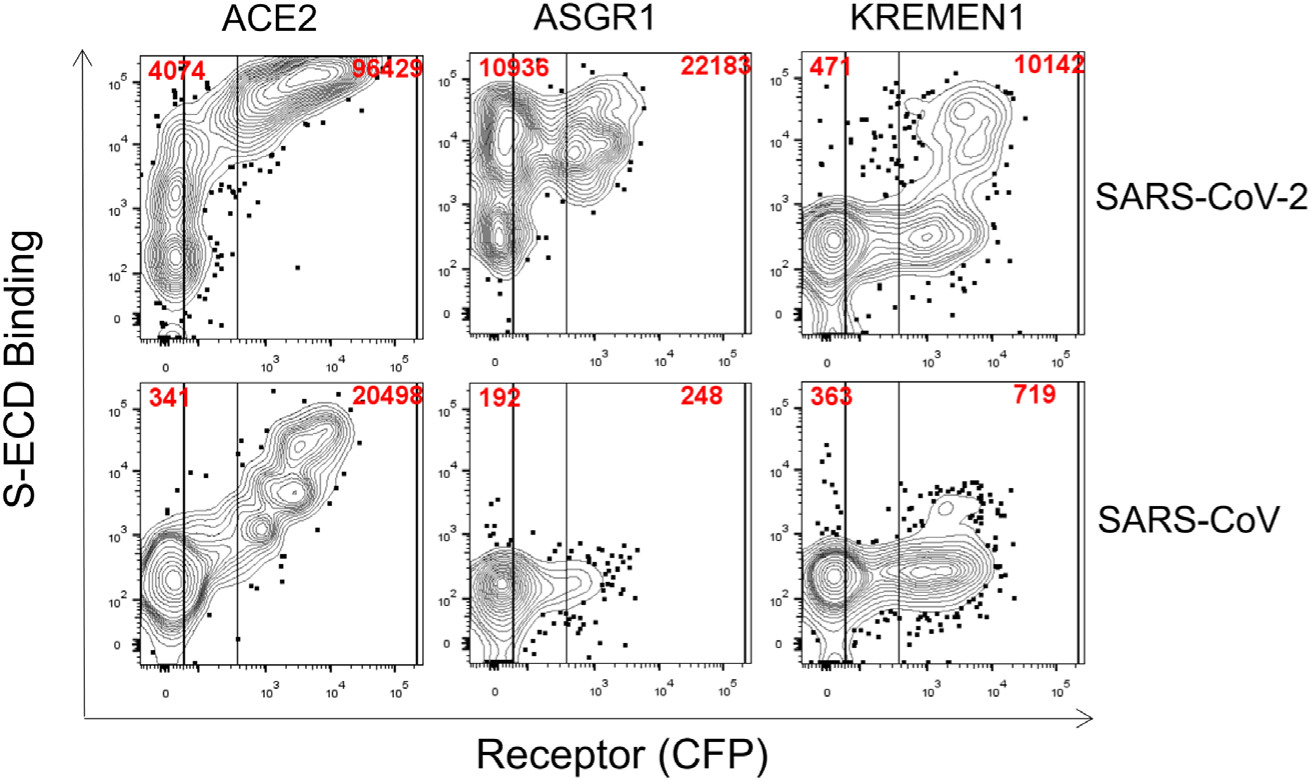
Binding of KREMEN1, ASGR1 and ACE2 with the S-ECD of SARS-CoV and SARS-CoV-2. KREMEN1, ASGR1 or ACE2 expressed 293e cells were incubated with S-ECD-hFc (10μg/ml final concentration) of SARS-CoV2 or SARS-CoV separately. S-ECD binding were measured by flow cytometry.

**Extended Data Fig.7.**
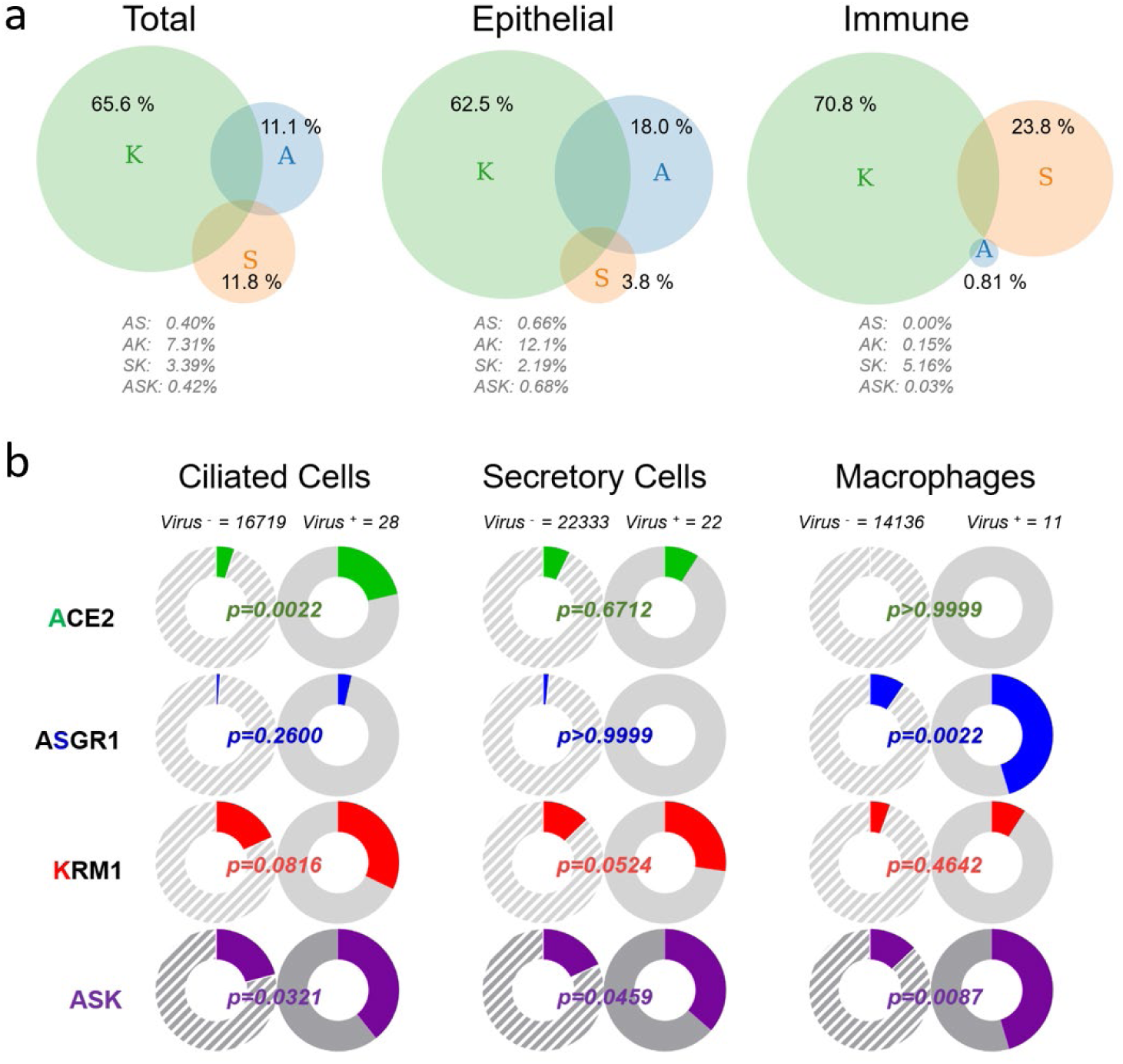
Distribution of ACE2, ASGR1 and KREMEN1 in ASK receptor positive cells and receptors correlation with virus susceptibility in the upper respiratory tract of patients with COVID-19. **a**, In ASK (<u>A</u>CE2/A<u>S</u>GR1/<u>K</u>REMEN1) receptor positive cells of total population, or epithelial and immune subpopulations of the upper respiratory tract with COVID-19, percentage of cells that expressing the indicated receptor(s) only were shown as venn diagram. **b**, In the epithelial ciliated and secretory cells, and immune macrophages of the upper respiratory tract with COVID-19, correlations of virus susceptibility with ASK receptors individually or in combination based on receptor positive cell percentage were determined. Virus positive and negative cell numbers were shown. p Values were calculated by Chi-square test.

**Extended Data Fig.8.**
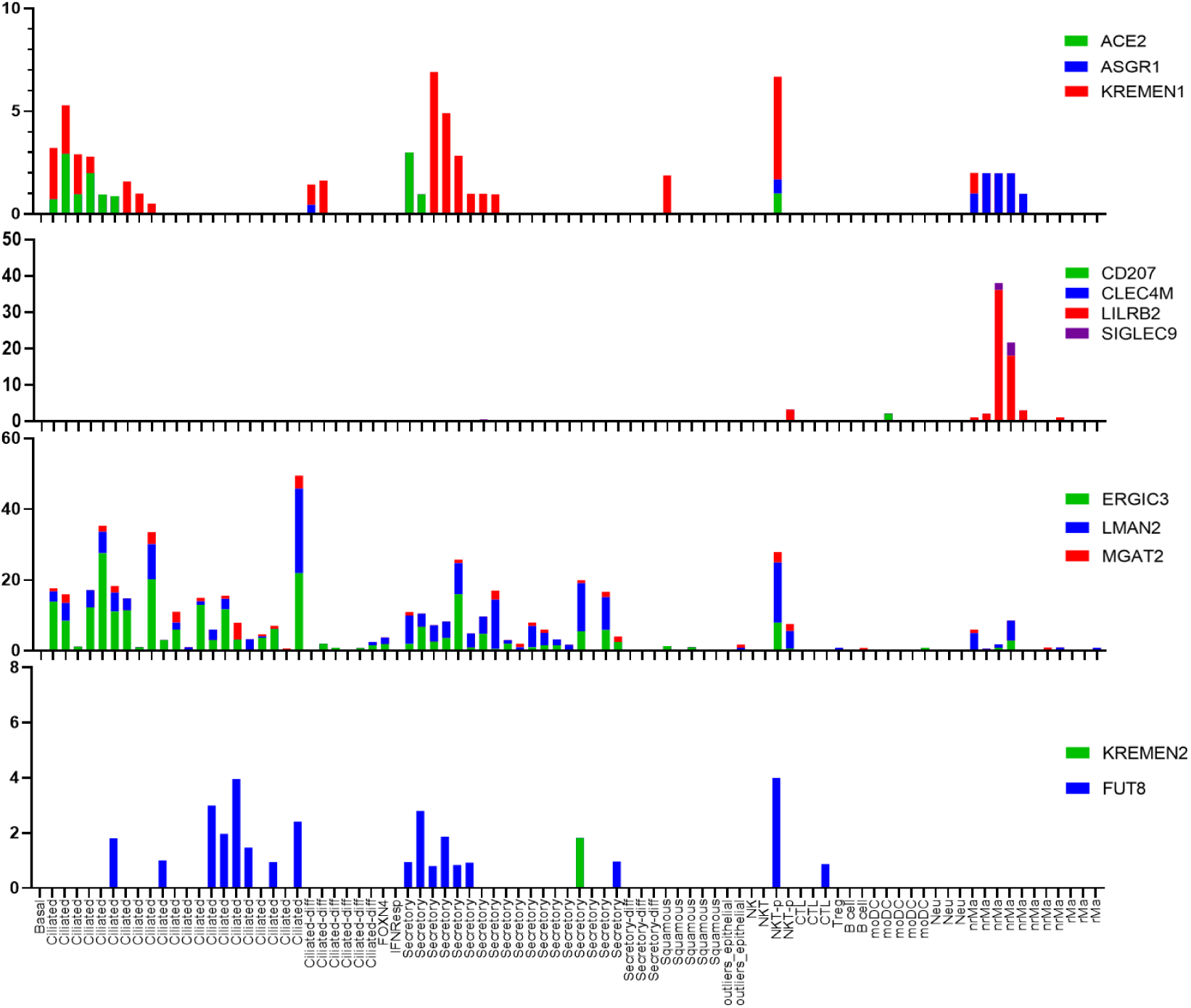
Expression of the twelve S receptors in SARS-CoV-2 positive cells from the upper respiratory tract with COVID-19.

**Extended Data Fig.9.**
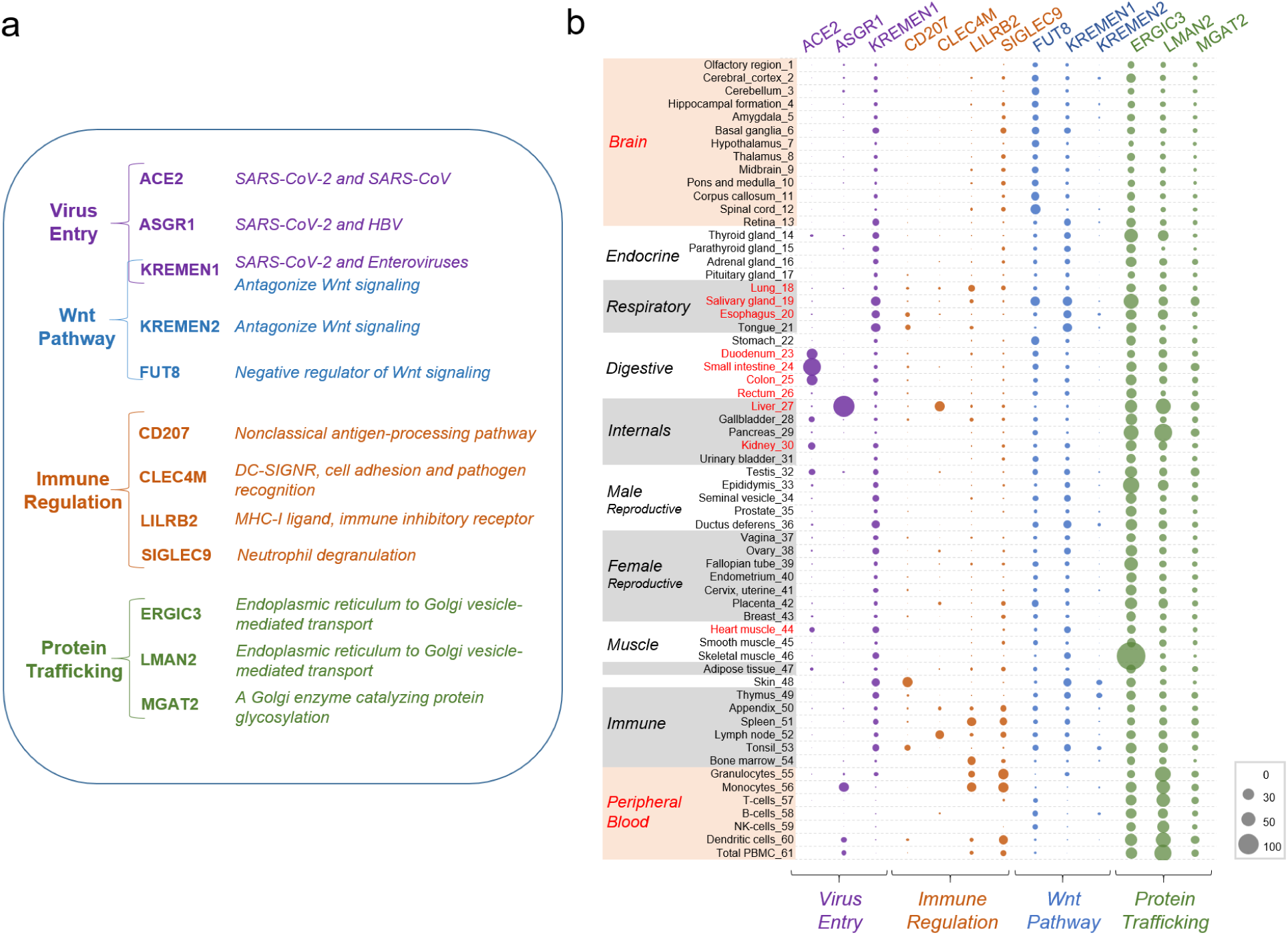
Expression pattern of SARS-CoV-2 receptors across human tissues. **a**, S receptors were classified according to their functions in virus entry, immune regulation, the Wnt pathway, and protein trafficking. **b**, Expression pattern of SARS-CoV-2 receptors across human tissues. mRNA expression levels of each receptor in human tissues were obtained from human protein Atlas. Tissues or organs that were identified as positive to SARS-CoV-2 are labeled red.

**Extended Data Fig.10.**
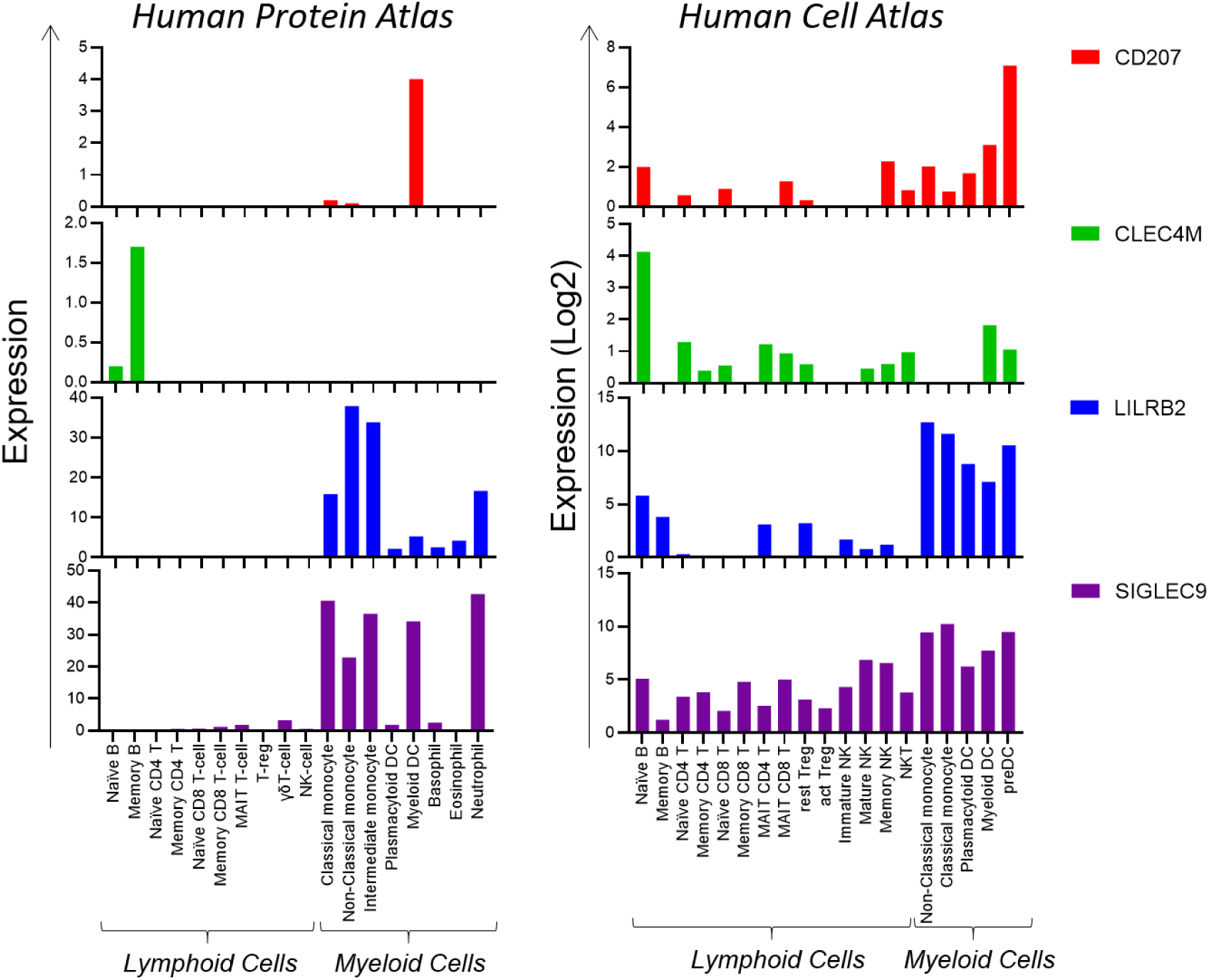
Expression pattern of CD207, CLEC4M, LILRB2 and SIGLEC9 in different cell populations of PBMCs. mRNA expression levels of indicated receptors in different cell population of PBMCs were derived from human Protein Atlas (https://www.proteinatlas.org/) and human Cell Atlas (http://immunecellatlas.net/) and shown as bar plot.

